# State-Dependent Dissociation of Shared Input and Directed Information Flow in the Visual Cortex

**DOI:** 10.64898/2026.01.22.701042

**Authors:** Yuxuan Xue, Mitchell Morton, Anirvan S. Nandy, Monika P. Jadi

## Abstract

Understanding how brain state and sensory input shape inter-laminar communication is essential for interpreting cortical population dynamics. Using laminar recordings in macaque V1, we apply reduced-rank regression to quantify low-dimensional predictive subspaces linking Input and Superficial layers. We find that both visual stimulation and internal state (eyes open vs. closed) modulate the structure and efficacy of these subspaces, but through distinct mechanisms. Visual input induces a directional, feedforward pattern from input to superficial layers, while wakefulness-related modulation enhances coordination more symmetrically. Delay analysis and network simulations confirm that structured, layer-specific inputs produce directional prediction, whereas global fluctuations yield undirected co-activation. Importantly, differential structure across layers predicts the emergence of communication asymmetry. These findings dissociate communication from shared modulation, providing a principled framework for interpreting inter-population correlations. Our results generalize to broader cortical circuits, offering insights into when population coupling reflects genuine information flow versus global state dynamics.

## INTRODUCTION

Animal behavior emerges from the interplay between external context (e.g., stimulus environment) and internal context (e.g., attention, arousal), which together shape perception, decision-making, and action [1–5]. These contextual states exert powerful influences on how neural circuits encode and process information. Understanding how such states modulate and interact within neural populations is essential for revealing core principles of brain function. A central unresolved question is how information is communicated between cortical populations in ways that both reflect and sustain these diverse behavioral states.

At the core of this question lies the concept of neural variability and correlation. Even under fixed stimulus conditions or identical behavioral tasks, cortical neurons exhibit substantial trial-to-trial variability in their responses [6–10]. This variability is not independent across neurons; rather, it is often shared in structured ways that can reveal underlying neural connectivity [8–14]. These trial-to-trial co-fluctuations, when analyzed at the population level, tend to occupy low-dimensional subspaces [1, 15–19], suggesting that only a few dominant patterns govern the majority of shared variance. Such population-level structure is often more informative than pairwise correlations, pointing to coordinated activity modes that are shaped by both anatomical constraints and functional demands.

Over the past decades, studies have examined spike-count correlations—the covariation in neuronal firing rates across repeated trials—in a variety of species, brain areas, and task conditions [8, 9, 11, 12, 14]. These studies have shown that correlation strength can be modulated by sensory input [6, 7, 13], attention [20–25], and state fluctuations [2, 4, 5, 8–10]. Importantly, even in the absence of external stimulation, the brain exhibits rich and structured spontaneous activity [1, 3, 26–29]. In many cases, spontaneous and stimulus-evoked activity share similar spatial and temporal patterns, suggesting that internal states may reflect latent models of the external world [30–34]. Recent work further shows that the baseline fluctuation defines a dominant axis that flexibly aligns with task-relevant signals, linking correlated variability to evoked function in behavior [35]. These insights strengthen our understanding of how contexts dynamically shape both spontaneous and evoked activity.

Despite these advances, a critical challenge remains: how to disentangle the distinct contributions of externally driven and internally generated sources of population correlation. Both can produce similar patterns of co-activation, yet they likely reflect fundamentally different mechanisms of circuit operation. For instance, strong correlations across populations of cortex during sensory stimulation may arise from feedforward transmission of sensory input, while similar correlations in darkness may reflect internally coordinated dynamics, such as ongoing cortical state fluctuations, top-down feedback, or global arousal signals [2, 4–11, 13]. Without a principled framework to dissociate these sources, correlation-based measures risk conflating communication with common input. This ambiguity is particularly relevant when interpreting correlations as evidence of neural communication. In addition to one population driving another, shared activity can also result from common inputs, oscillatory coupling, or broader brain-state modulations that affect multiple regions simultaneously [2, 4, 5, 7–11, 13, 20, 23, 36–38]. Thus, we must ask: when does correlation reflect direct information flow, and when is it merely a signature of shared modulation?

To address this, recent studies have leveraged a Reduced-rank Regression (RRR) framework based on predictive modeling, which allows us to move beyond simple measures of correlation strength [25, 39–42]. Specifically, these approaches use predictive subspace analysis, which characterizes the dominant low-dimensional patterns of activity in one population that are most informative about activity in another. Unlike traditional PCA or FA, which reduce dimensionality within a single population, such approaches apply reduced-rank regression across populations, capturing the predictive structure that links them.

In this study, we extend this approach to detect asymmetric relationships and distinguish feedforward-driven co-fluctuations from symmetric, spontaneous ones. Importantly, we utilize the directionality of prediction—how well one population can predict another, and whether this prediction is asymmetric—as a metric of information flow. Our study focuses on the primary visual cortex (V1) in macaques, where cortical layers are anatomically and functionally well-characterized. V1 receives robust bottom-up input and has been a model system for understanding feedforward processing [30, 34, 43–45]. Yet even in V1, spontaneous activity exhibits remarkable structure and coordination across layers [4, 8–10, 42, 46, 47]. Previous studies have demonstrated that inter-population communication subspaces are low-dimensional, including between Input and Superficial layers in V1, and are modulated by stimulus conditions [25, 40–42, 48]. What remains unclear is whether subspace structure changes across different contextual states (Fig. 1A), as well as the extent to which observed correlations reflect true information transmission. This leads us to several motivating questions: Can we dissociate visually driven and spontaneous sources of population correlation in V1? Does inter-laminar correlation reflect true directional communication, or shared internal dynamics? How do changes in external input and internal state reshape the functional architecture of cortical networks?

**Fig 1.**
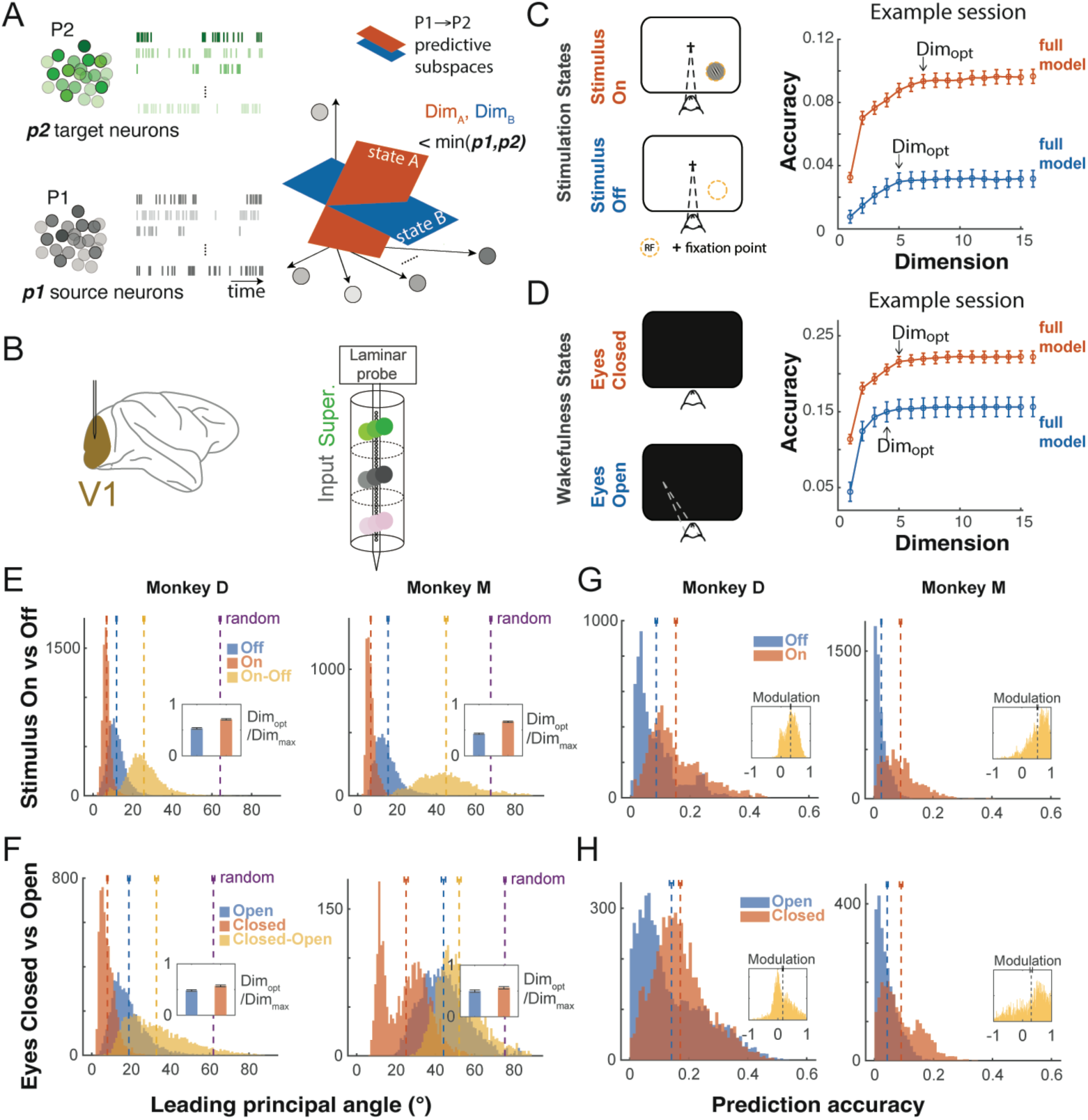
Modulation of inter-laminar predictive subspace by sensory and wakefulness states in macaque V1. (A) Schematic of a low-dimensional predictive subspace. The activity of target population (green) is projected onto a subspace of the activity space of source neurons (grey). Schematic shows a hypothetical modulation of this subspace by contexts such as wakefulness and presence of sensory inputs. (B) Illustration of electrophysiological recordings. The considered neurons include single units and multi-units. Layers were identified using current-source-density (CSD) mapping (see Methods). (C-D) Reduced-rank regression (RRR) using input activity to predict superficial activity. Examples are shown for monkey D. The error bars on the prediction accuracy indicate the standard error across multiple draws of trials and the corresponding cross-validation folds. Full model prediction is obtained from Ridge regression. (E) The distribution of the leading principal angle (LPA) during visual stimulation. The dashed lines show the mean LPA with error bars indicating the 95% confidence intervals (CIs) of the mean, obtained by repeatedly subsampling from the data. LPA is obtained from comparing subspaces across stimulus conditions (on vs. off; yellow) and within stimulus conditions: off (blue) or on (orange). The purple line indicates LPA calculated from random subspaces with the same dimensions as stimulus on vs. off. Inset: the fractional optimal dimensions. (F) The distribution of LPA during spontaneous sessions. LPA is obtained from comparing subspaces across wakefulness conditions (eyes open vs. closed; yellow) and within wakefulness conditions: eyes open (blue) or eyes closed (orange). (G) The distribution of prediction accuracy during visual stimulation. The dashed lines show the mean accuracy with error bars indicating the 95% CIs of the mean. Orange: stimulus on; Blue: stimulus off. Inset: the modulation index (MI; see Methods) distribution for all pairs of compared samples. (H) The distribution of prediction accuracy during spontaneous sessions. Orange: eyes closed; Blue: eyes open.

To answer these questions, we combine laminar recordings in macaque V1 [9, 24, 42] with a predictive modeling framework that quantifies directed and undirected population coupling. We examine how patterns of inter-laminar prediction change across behaviorally distinct conditions and test whether directional asymmetry is preserved or disrupted. Further, we simulate simplified models of cortical circuits to test whether different sources of co-fluctuation produce distinguishable signatures in prediction metrics.

While our primary goal is to distinguish visually driven from spontaneous co-fluctuations, our broader aim is to develop analytical tools for interpreting population correlations in terms of underlying circuit mechanisms. This work builds on prior efforts to define communication subspaces [40–42], but shifts the focus from decoding or representational content to the structure of population-level interactions. Ultimately, we argue that predictive subspace analysis—especially when combined with careful state manipulation and anatomical resolution—provides a powerful lens through which to view functional connectivity in the cortex.

## RESULTS

We aimed to investigate the structure and efficacy of the inter-laminar subspace in V1 and how it is modulated by sensory input and wakefulness state. We specifically focused on the Input and Superficial layers, which constitute a canonical information processing pathway in the sensory neocortex [34, 49]. To this end, we analyzed neural recordings from macaque V1, where electrophysiological signals were simultaneously recorded across cortical layers (Fig. 1B) over multiple sessions in two animals, with concurrent eye tracking. We examined both stimulus-evoked and spontaneous activity via trial-by-trial variability. Grating stimuli were repeatedly presented at the receptive field (RF) for 100 ms, followed by a 200–250 ms interstimulus interval. For each trial, spikes were counted in 100-ms windows before and after stimulus onset, corresponding to different sensory processing states. We also assessed spontaneous activity during periods of darkness, with spike counts binned in 500-ms windows, while the animals’ eyes were either open or closed, corresponding to distinct wakefulness states. Inter-laminar coordination between the Input and Superficial layers was quantified using Reduced Rank Regression (RRR; see Methods), during which we implemented Ridge regression to prevent overfitting (Fig. S1). This procedure identified low-dimensional predictive subspaces for linear models mapping activity from input to superficial populations.

### V1 inter-laminar predictive subspace is modulated by sensory and wakefulness states

We found that the predictive subspace was low-dimensional and modulated by both sensory and wakefulness-related inputs. In all four conditions, prediction accuracy reached a plateau at a reduced dimensionality, beyond which additional predictive dimensions no longer improved performance (Fig. 1C, D). The predictive subspace was defined as the regressor at this optimal dimensionality, marking the most informative correlation structure between the two layers.

To compare subspace structure across conditions, we computed the leading principal angle (LPA; see Methods) between predictive subspaces obtained under different states. For consistency, the same number of Input and Superficial neurons was sampled across sessions within each monkey. The mean LPA between stimulus-on and stimulus-off conditions was significantly larger than the within-condition variability (Fig. 1E), indicating that visual input induces a consistent reorganization of the inter-laminar subspace. These LPAs were consistently smaller than those between randomly oriented subspaces of the same dimensionality, suggesting the presence of structured, non-random connectivity. A similar pattern was observed when comparing eyes-open and eyes-closed states (Fig. 1F), indicating that wakefulness also modulates inter-laminar coordination.

In addition to these structural subspace changes, prediction accuracy—reflecting the strength of inter-laminar correlation—was enhanced under both visual stimulation (Fig. 1G) and eye-closed conditions (Fig. 1H). Together, these results demonstrate that both external sensory input and wakefulness state shape the structure and efficacy of inter-laminar functional connectivity in V1.

### Predictive subspace is modulated by sensory and wakefulness states in distinct ways

Although both sensory and wakefulness-related inputs modulate inter-laminar coordination, we hypothesized that they do so via distinct mechanisms, potentially arising from different sources of drive to this pathway. To test this, we evaluated the directionality of coordinated activity between neuronal populations in the two layers by comparing prediction performance under two modeling frameworks–directed and undirected. In the directed model, neurons in the Input layer served as the source population, with the superficial layer as the target (Fig. 2A, left). In contrast, the undirected model involved randomly assigning source and target populations from a pooled set of neurons drawn from both layers (Fig. 2A, right). For both models, we applied the same analysis pipeline used in Fig. 1G, H to compute prediction accuracy and derived the corresponding modulation index (MI) (Fig. 2B).

**Fig 2.**
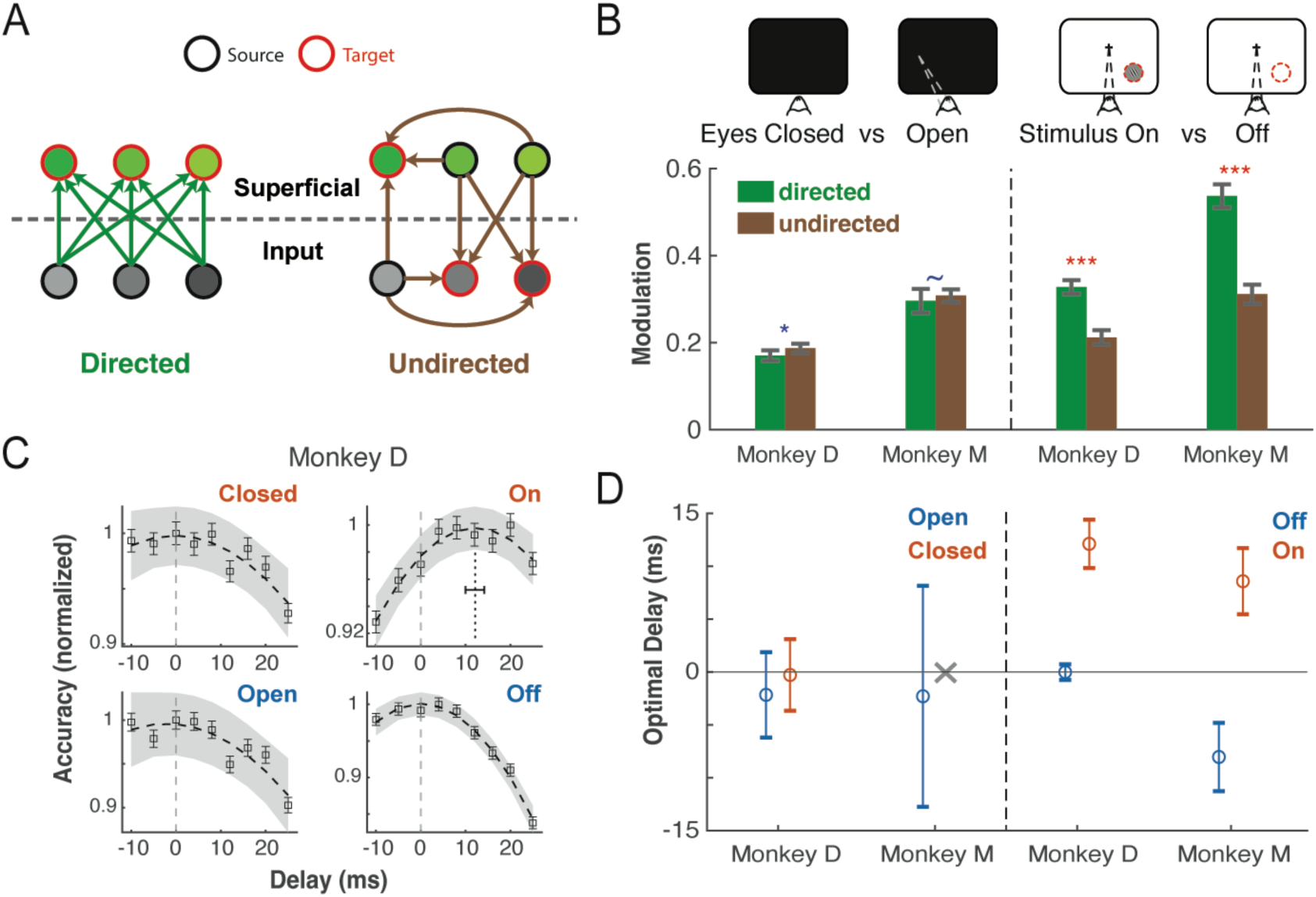
Two mechanisms for modulating the predictive accuracy across Input and Superficial layers. (A) Schematic of two possible prediction patterns among Input and Superficial neurons. (B) Prediction accuracy modulation index (MI) for directed vs. undirected models during spontaneous (left panel) and visual stimulation (right panel) conditions. Green: directed model, same as Fig. 1G, H insets; Brown: undirected model, where source and target populations were randomly sampled from all neurons pooled together. Error bars indicate the 95% CIs of the mean. Significance was assessed using two-sample t-tests. (C) Delay analysis of the directed model in monkey D. The target activity of the superficial population is shifted in the range -10 ms to 25 ms w.r.t. Input layer activity. The accuracy is normalized, then fitted to a quadratic function of the delay time. Yellow areas represent the 95% CIs of the normalized efficacy. The vertical dotted line in stimulus-on condition shows the symmetry axis (optimal delay), along with the standard error. (D) The estimate on optimal delays for all four conditions in two monkeys. The error bars indicate the standard error. The cross mark indicates a poor fit (no clear dependence of accuracy on the delay time).

Across both monkeys, the modulation by sensory input was significantly higher for the directed model compared to the undirected model (Fig. 2B, right), indicating that the predictive subspace is preferentially enhanced in the feedforward direction—from input to superficial layers—rather than in an arbitrary or bidirectional manner. In contrast, under different wakefulness states, this directionality was not observed: the MI for the directed model was not significantly different from, and in one of the two monkeys even lower than, that of the undirected model (Fig. 2B, left). This suggests that wakefulness modulates the efficacy of cross-layer coordination in a more global, non-directed fashion, enhancing population interactions across layers without imposing a specific source–target structure. To further examine the directionality of information flow, we performed a delay analysis (see Methods) on the directed model. We identified the optimal time lag at which Input layer activity best predicted superficial layer responses (Fig. 2C, S2). Under visual stimulation, predictive models were fitted when aligning activity to response onset for each layer, ensuring that any observed delays in predictive accuracy could not be attributed to latency differences. The optimal delay consistently occurred at a positive value, indicating that Input layer activity precedes and predicts superficial layer activity. This trend was robust across both monkeys (Fig. 2D). In contrast, the other three conditions—stimulus off, eyes open, and eyes closed—did not exhibit a consistent non-zero delay. These findings support a causal interpretation: visual stimulation activates the Input layer, which in turn drives superficial layer activity through inter-laminar interactions. During spontaneous activity, however, the interaction between layers appears more symmetric and complex, consistent with the absence of directed enhancement observed in the MI analysis.

### Simulating a two-layer RNN with layer-specific external inputs

To gain further insights into the mechanisms underlying the directional modulation in V1, we simulated a simplified two-layer recurrent neural network (RNN) designed to replicate key features of the changing predictive subspace structure and efficacy observed in the data. The network consisted of two fully connected layers, each comprising 100 units with uniform intra-layer connectivity. The feedforward connections were modeled as low-rank, randomly positive matrices, while feedback from the superficial layer to the Input layer was set to zero for simplicity (Fig. 3A; see Methods). Neuronal activity *x*_*i*_ evolved according to a standard recurrent update rule

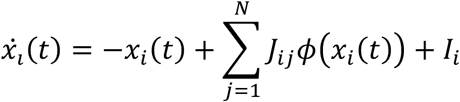

**Fig. 3.**
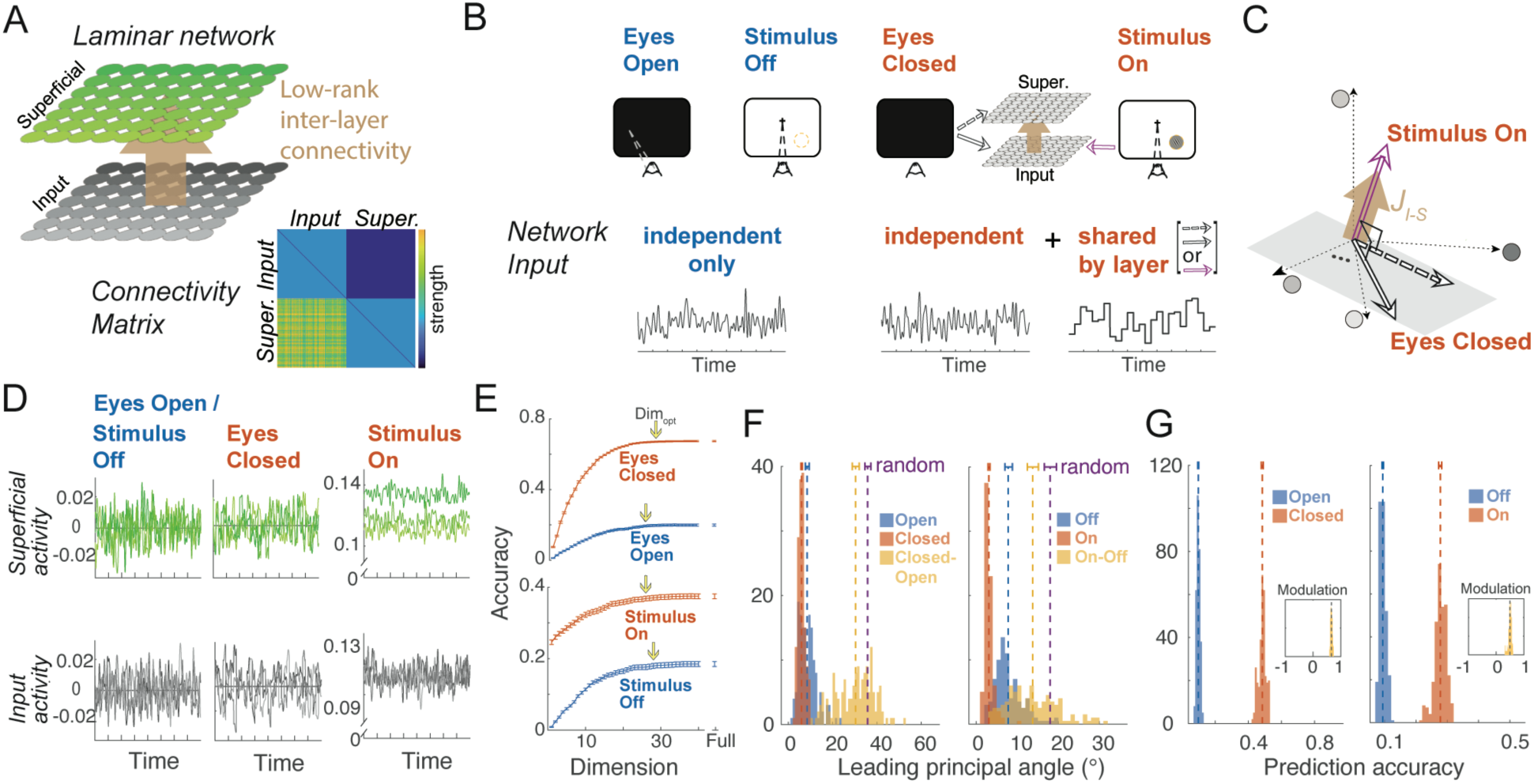
Modeling directed and undirected mechanisms of predictive subspace modulation in a two-layer recurrent neural network. (A) Schematic of the two-layer network model. Each layer is fully recurrently connected, with low-rank feedforward connectivity from the input to the superficial layer and no feedback. (B) External input structures under four conditions. In the resting state (eyes open/stimulus off), neurons receive independent Gaussian noise. In the eyes-closed condition, an additional shared fluctuation targets both layers. In the stimulus-on condition, an additional shared fluctuation specifically targets the Input layer. (C) Under stimulus-on conditions, the direction of the additional input at any time aligns with the dominant modes of the feedforward connectivity matrix J_I−S_ (brown arrow). The additional shared inputs under eyes-closed conditions (solid: Input layer; dashed: superficial layer) are orthogonal to the dominant modes. (D) Simulated activity *x*_*i*_ from example neurons (three in each layer) under resting state (left), eyes-closed (mid), and stimulus-on (right) conditions. (E) RRR results for predicting superficial layer activity from Input layer activity. Example shown for a 40-neuron source and 40-neuron target. The optimal dimensionality is marked where prediction accuracy saturates. (F) The distribution of LPA. Left: eyes-closed vs. resting state; right: stimulus-on vs. resting state. (G) The distribution of prediction accuracy. Left: eyes-closed vs. resting state; right: stimulus-on vs. resting state. Insets: MI of prediction accuracy induced by the external inputs of either condition.

where *ϕ* is a nonlinear current-to-rate function, and *J* is the connectivity matrix.

We simulated three input conditions: a baseline input representing either the eyes-open or stimulus-off condition, an additional input corresponding to eyes-closed, and another corresponding to stimulus-on. At baseline, both layers received only independent Gaussian noise (Fig. 3B). The eyes-closed condition introduced shared input fluctuations targeting both layers, emulating spontaneous cortical drive from upstream areas. In contrast, visual stimulation introduced structured inputs targeting only the Input layer. To differentiate the latter two conditions, the stimulus input was designed to align with the dominant modes of the inter-layer connectivity matrix *J*_*I*–*S*_, maximizing its impact on downstream neurons. Conversely, spontaneous input was constructed to be orthogonal to the dominant modes, thus minimizing directionality (Fig. 3C).

In both layers, simulated activity exhibited fluctuations around different mean levels (Fig. 3D). As designed, visual input elevated the mean activity of both layers, consistent with empirical data. We then applied the same RRR-based directed model analysis to the simulated data (see Methods), extracting predictive subspaces, leading principal angles (LPA), and prediction accuracy. The results showed that coordinated inter-layer activity remained low-dimensional (Fig. 3E), and that both spontaneous and visually driven inputs modulated the subspace structure similarly (Fig. 3F). Correspondingly, prediction accuracy increased under both conditions, indicating enhanced subspace efficacy (Fig. 3G). Overall, the simulated network reproduced the key features observed in the empirical data, demonstrating that structured input—whether stimulus-driven or during wakefulness—modulates a low-dimensional, functionally meaningful mapping between Input and Superficial layers.

### The relative activity structure across layers determines the directionality of inter-layer predictive performance

Using the simulated network, we aimed to better understand what drives the directionality of modulation induced by sensory and wakefulness inputs. While the structured nature of external input and its layer-specific targeting are essential features of the model, they do not necessarily guarantee directional predictive performance. Previous work has shown that input neurons driven by the lateral geniculate nucleus (LGN) exhibit more independent activity, while superficial neurons show more coordinated activity during visual stimulation [10]. Yet little is known about how such laminar differences translate into directional communication, nor about how spontaneous activity shapes interactions across layers. We hypothesized that the relative activity structure across layers would impact the directionality of inter-laminar information flow. To test this, we systematically varied the amplitude of independent Gaussian noise input across the two layers to assess its effect on directionality.

We first replicated the empirical result that communication during visual stimulation is directed from the input to the superficial layer, whereas spontaneous co-fluctuations in the eyes-closed state are largely undirected (Fig. 4A). Building on this, we found that the ratio of independent noise amplitudes between Input and Superficial layers plays a critical role in determining whether modulation is directional during visual stimulation. Specifically, only when the Input layer shows stronger independent fluctuations does the sensory-driven modulation become directed from input to superficial layer (Fig. 4B). In contrast, modulation induced by the eyes-closed state—where activity is dominated by shared fluctuations—remained largely undirected, regardless of the ratio of independent noise to each layer (Fig. 4B). These results suggest that the layer-specific targeting (Fig. 3B) alone is insufficient to capture the distinction between stimulated and spontaneous conditions. In the stimulated case, directional communication requires lower independent noise in the superficial layer, whereas spontaneous activity does not engage such a mechanism.

**Fig. 4.**
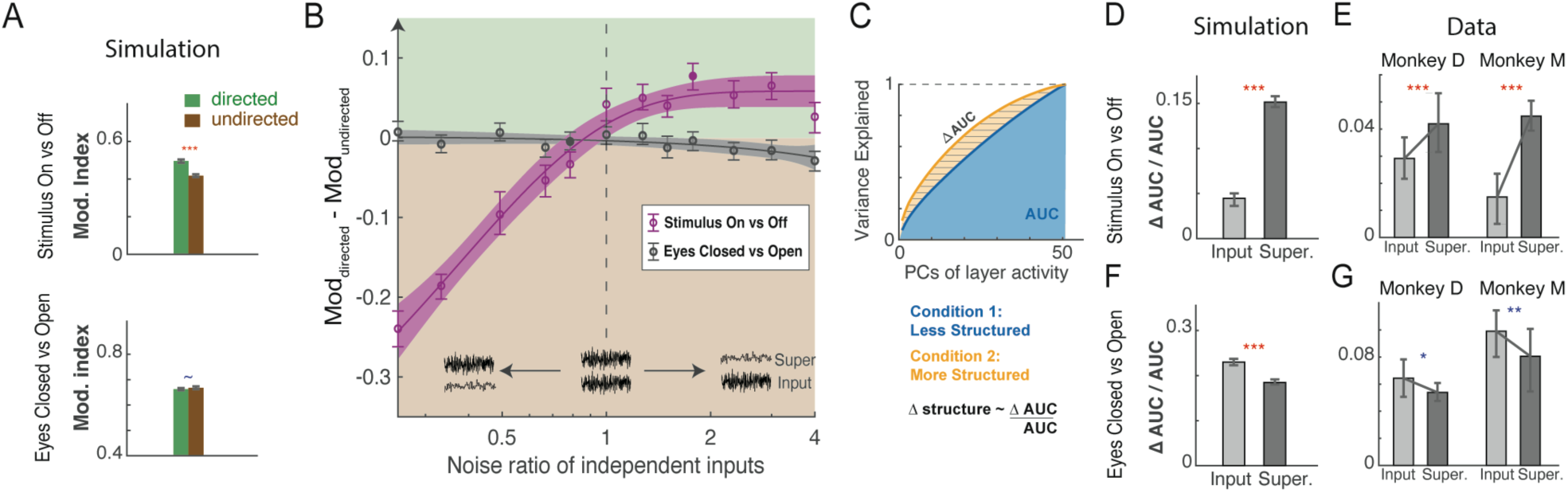
The relationship between relative activity structure and the directionality of inter-layer prediction. (A) Prediction accuracy modulation index (MI) for directed vs. undirected RRR models during stimulated (top) and spontaneous (bottom) periods in the simulated network model. Error bars indicate the 95% CIs of the mean. Significance was assessed using two-sample t-tests. (B) Directionality of inter-layer activity—defined as the difference in MI between the directed and undirected models—varies systematically with the ratio of Gaussian noise in the independent inputs of Input and Superficial layers. Green/Tan shaded areas indicate regions where the simulated activity is considered layer-directed/undirected. The x-axis is plotted on a logarithmic scale. Each circle represents a data point, with error bars indicating the 95% CIs of the mean. Solid circles correspond to conditions analyzed in (A), (D) and (F). Colored curves represent exponential fits to the data, with shaded regions indicating the 95% CIs of the fit. Positive y-values indicate that activity is preferentially directed from input to superficial layers. Such directional co-activity emerges only when the ratio exceeds 1 for the stimulated condition (top right of the purple curve). (C) Definition of population activity structure. For a given population, the area under the curve (AUC) of the cumulative-variance-explained across principal components quantifies the low-dimensional dominance. A larger AUC (red area compared to blue) indicates a higher level of structured population activity. (D) The increase of structured activity in Input and Superficial layers under stimulus on vs. off conditions. Error bars indicate the 95% CIs of the mean. Significance was assessed using two-sample t-tests (also applied in E-G). (E) Same increase shown for empirical data from two monkeys. (F) The increase of structured activity in Input and Superficial layers under eyes closed vs. open conditions. (G) Same increase shown for empirical data from two monkeys.

In our model, lower independent noise within a layer is associated with more structured, lower-dimensional activity—reflecting greater dependence among neurons. Such structured activity may arise in the superficial layer through local recurrent processing of input-layer signals. To quantify this effect, we introduced a metric for activity structure based on principal component analysis (PCA): for each population, we computed the cumulative variance explained by the ordered principal components and defined the area under the curve (AUC) as the level of activity structure (Fig. 4C; see Methods). An upward shift in the variance curve indicates greater low-dimensional dominance, and therefore more structured patterns in the population activity. We then quantified the modulation of activity structure within each layer by measuring the change in AUC under different experimental conditions. We observed that, for sensory-driven modulation that is directed from input to superficial layer, visual stimulation selectively enhanced the activity structure of the superficial layer more strongly than that of the Input layer (Fig. 4D). This relative structure is consistent with empirical data in both monkeys (Fig. 4E). In contrast, neural recordings from the spontaneous condition showed the opposite trend: the Input layer exhibited greater increase of structure due to eyes closing (Fig. 4G). Our simulation model also successfully reproduced the input-superficial structure (Fig. 4F). While the simulated modulation maintains undirected for any parameters, the input-superficial structure increase can be reproduced only within a specific range, particularly when the noise ratio is less than 1 (Fig. S3).

Together, these results demonstrate that visual stimulation induces distinct levels of activity structure across layers, and this layer asymmetry supports directional predictive performance from input to superficial layers. Wakefulness-related inputs, on the other hand, yield the opposite pattern of layer-specific organization, but such a difference does not translate into directional modulation. Instead, spontaneous interactions remain largely undirected, regardless of structural differences between layers.

## DISCUSSION

Our study reveals that inter-laminar co-fluctuations in the primary visual cortex arise from two distinct mechanisms: one driven by sensory input and the other by internally generated dynamics associated with wakeful rest, both of which modulate the structure of the predictive subspace. Under visual stimulation, predictive performance consistently favors a directed flow of information from input to superficial layers, aligning with known anatomical feedforward pathways engaged during sensory processing [30, 34, 41–44, 48]. In contrast, under spontaneous conditions, this directionality diminishes, and co-fluctuations become more symmetric, with an undirected model more effectively capturing the observed interactions.

These empirical findings are further supported by simulations, which demonstrate that increasing the relative activity structure of the superficial layer enhances its predictability from the Input layer—reproducing the directionality observed during sensory-driven activity. This modeling framework suggests that the Input layer, being less structured, is capable of representing a broader range of stimuli [9, 10, 34, 42, 45], while the superficial layers, shaped by local recurrence and top-down feedback, promote reliability and structured encoding—facilitating communication within recurrent and feedback loops [24, 38, 43, 50]. In this view, directionality in predictive subspace models naturally emerges from differences in representational structure across layers, offering a mechanistic explanation for the asymmetric inter-laminar dynamics observed under sensory stimulation.

### Dissociating Visually Driven and Spontaneous Inter-laminar Co-fluctuations

Building on the observed differences in predictive model performance across task contexts, we interpret these results as evidence for two distinct mechanisms of inter-laminar coordination. During visual stimulation, the enhanced predictability from input to superficial layers reflects causally structured, feedforward communication, consistent with the notion that visual input primarily targets the Input layer and propagates through local circuits to influence superficial activity. In contrast, under spontaneous conditions, co-fluctuations become more symmetric, suggesting a regime dominated by shared input or globally coordinated internal dynamics, rather than directed transmission.

Spontaneous and evoked mechanisms likely coexist in naturalistic scenarios, and even during passive viewing conditions, internally generated fluctuations are superimposed on stimulus-driven responses [1, 3, 8–10, 28, 32, 33]. This view is supported by recent insights into the predictive structure of spontaneous activity, which suggest that spontaneous fluctuations are not random noise but instead reflect organized internal dynamics [1, 3, 32, 35, 51–53]. Indeed, a growing body of literature demonstrates remarkable similarity between spontaneous and sensory-evoked activity, both in spatial and temporal structure, across the visual cortex [35, 51, 54, 55], auditory cortex [52, 56], and hippocampus [57–59], and across a variety of species and stimulus types [1, 35, 51, 54]. These findings reinforce the idea that spontaneous activity encodes functionally relevant patterns that mirror and perhaps anticipate sensory input.

Given the overlap between spontaneous and evoked activity, dissociating their circuit-level underpinnings requires an investigative framework beyond co-fluctuations. Our framework addresses this by contrasting extreme conditions of external input structures and manipulating the relative structure of activity across layers. The Input layer with its higher-dimensional structure naturally supports more diverse representations and is more likely to drive downstream activity. When such activity targets less structured, more recurrent or integrative populations—like the superficial layer—this asymmetry becomes functionally meaningful and measurable. Thus, predictive asymmetry emerges not solely from anatomical hierarchy, but from how information is encoded, propagated, and transformed across structured populations.

This modeling principle—tuning structural asymmetry to reveal functional directionality—can be extended to other systems beyond V1. For instance, it may help dissociate feedforward and feedback interactions in higher sensory areas, or tease apart internally generated sequences versus sensory-evoked responses in hippocampus or prefrontal cortex. More broadly, it offers a generalizable framework for probing how circuit architecture and representational geometry jointly shape the flow of information in the brain.

### Rethinking Correlation as a Marker of Communication

While functional correlations across cortical layers are often interpreted as evidence of communication, our results suggest that such correlations can arise from fundamentally different mechanisms—either through feedforward transmission of sensory input, or from shared, spontaneous fluctuations. This distinction becomes especially important in non-stimulus-driven conditions, where apparent inter-laminar coupling may reflect global brain states rather than direct information transfer.

Our findings do not contradict prior work identifying communication subspaces during visual tasks with well-controlled input [25, 40–42]; indeed, in such contexts, feedforward drive dominates, and communication is more prominent. Rather, we emphasize that the nature of inter-laminar correlation may vary depending on the contextual state of the animal. Spontaneous fluctuations can mimic or mask directed communication if not properly accounted for. This concern aligns with interest in using Bayesian methods to infer latent brain states from neural data and underscores the importance of considering state-dependent dynamics when interpreting population coupling [15, 19, 60, 61].

Temporal information in our recordings further supports the distinction between these mechanisms. Even after aligning neural activity to the onset of visual responses across layers, we observe that predictive accuracy from input to superficial populations peaks at a delay (Fig. 2C)—consistent with local recurrent processing in the superficial layers. This suggests that Input and Superficial layers are not merely passively co-active, but operate in sequence, with distinct temporal dynamics. Such evidence supports the view of communication as a stage-wise process of information transformation, rather than merely shared activation. The anatomical grounding afforded by our CSD-based identification of laminar boundaries strengthens this interpretation by ensuring accurate assignment of recorded units to cortical layers [46].

### Predictive Subspaces Reflect Functional Connectivity

We use predictive subspace analysis to characterize the functional relationship between Input and Superficial layers in V1. Rather than reducing dimensionality within each layer independently (e.g., PCA or FA), our method applies reduced-rank regression across populations, targeting the cross-layer predictive structure. This design ensures that the extracted subspace captures inter-population prediction—a signature of functional connectivity, rather than internal redundancy. Prior work has shown that these predictive dimensions are distinct from the principal axes within each population alone, reinforcing the idea that this method isolates a population interaction space, not necessarily a low-dimensional representation of activity within a single layer [41, 42].

This distinction is important because the low-dimensional nature of single-population neural activity is well established [15, 17, 19], whereas investigations into the nature of the low dimensionality of inter-laminar predictive subspaces are at a nascent stage. To understand whether these subspaces reflect genuine features of cortical circuitry, we examined the orientation of the predictive subspace across contextual states and found consistent shifts (Fig. 1E,F). These rotations indicate genuine changes in the underlying connectivity patterns. The dimensionality of the predictive subspace also appeared to vary systematically across conditions (Fig. 1E,F, insets). However, we remain cautious in interpreting the dimensionality changes as robust subspace properties due to the limited number of target neurons that makes optimal dimension estimates discrete and noisy. Furthermore, the optimal dimension is sensitive to other factors such as recording noise, sampling variability, and model hyperparameters [16, 19, 62]. To reduce potential biases, we fixed all model parameters and avoided neuron selection or tuning for performance, thereby ensuring that the observed subspace rotations reflect real structural shifts rather than dataset-specific artifacts. Given these limitations, we focus our interpretation on the reliable changes in orientation and defer conclusions about dimensionality until denser recordings become available.

To further support our conclusion that predictive subspaces reflect meaningful network-level structure, we performed simulations in which we manipulated the asymmetry of activity and independent noise across populations (Fig. 4B). These synthetic datasets showed that the functional role of the predictive subspace—whether it supports directional flow of information or more symmetric co-activation—depends critically on underlying network mechanisms. In other words, the subspace does not simply reflect signal complexity or shared variance; instead, it adapts to the distinct architecture of input-superficial communication.

Taken together, these findings suggest that predictive subspaces reveal circuit-level properties of cortical organization. Rather than echoing the internal redundancy of single populations or emphasizing absolute dimensionality, the subspace captures how populations functionally interact across layers. While its orientation shifts with contextual demands, its functional role—as either a communication pathway or a coordination scaffold—is shaped more by network architecture than by context.

### Modeling Inter-laminar Interaction: Insights and Limitations

Our modeling framework was designed to isolate and illustrate how different types of input drive distinct patterns of inter-laminar co-fluctuation. Specifically, we simulated two input regimes: one that selectively activates the Input layer and the input-to-superficial pathway, and another that drives both layers in parallel, designed such that the drive does not directly engage the inter-layer pathway (Fig. 3C). These two conditions serve as idealized representations of visually-driven and spontaneous-type co-fluctuations, respectively. Despite the model’s simplicity, it replicates key signatures observed in our data and supports our interpretation that the origin of co-fluctuation—whether causal or shared— can be inferred from the structure and directionality of predictive accuracy.

While the model includes no explicit feedback pathways, these pathways may play a significant role in spontaneous co-fluctuation [34, 36, 63]. For example, signals from higher areas such as V2 may project back to both deep and Input layers of V1 [64–66], potentially synchronizing activity across layers without driving feedforward computation. Similarly, local feedback within V1 (e.g., from deep to superficial layers) could reinforce co-activation patterns, particularly in the absence of visual input [67, 68]. These mechanisms may contribute to the correlation structure we observe during spontaneous activity.

Although such local feedback is difficult to isolate experimentally, computational approaches could in principle begin to disentangle its contributions. For instance, future models incorporating layer-specific populations and recurrent motifs could help distinguish whether observed co-fluctuations are better explained by shared upstream input, feedback loops, or a combination of both. In the current study, however, our focus is not on resolving these mechanisms in full detail, but rather on establishing how predictive subspace structure reflects functional coordination across layers, irrespective of precise circuit origin.

Our model lacks details such as the spatial architecture of recurrence [69], layer-specific cell types [43, 70], cell-type specific firing [71, 72] and synaptic physiology [73]. Still, even in its current form, the model offers a compelling proof of principle: that co-fluctuation patterns can reveal not just whether two populations are correlated, but how information flows between them—or fails to. Biologically enriched modeling could serve as a critical next step for investigating how specific anatomical features—such as feedback, cell-type interaction, or subcortical inputs—give rise to the observed co-fluctuations.

### Future Directions and Broader Implications

This study provides a foundation for interpreting inter-laminar population coupling as a marker of distinct sources of co-fluctuation—sensory-driven versus spontaneous—and for linking these patterns to mechanisms of neural communication. While our analysis focused on V1 laminar populations, population-level correlations have been widely studied in areas such as the hippocampus [74, 75], prefrontal cortex [76, 77], and higher visual cortical areas [20, 24, 78, 79], where they are often taken as signatures of shared coding, network state, or functional connectivity. In these systems, correlated variability has been linked to memory consolidation [75, 80], decision-making [7, 79], attention [20–22], and internal dynamics such as replay or persistent activity [1, 26, 81], and more broadly to flexible population codes that bridge spontaneous and evoked activity [35].

However, population correlation is an inherently ambiguous measure: it may reflect direct communication (i.e., information flow), but also common input or global brain states that simultaneously modulate multiple populations. For example, hippocampal theta and ripple events can drive synchrony across layers and subfields without implying directed interaction [80, 82]. Similarly, in prefrontal cortex, correlations often shift with behavioral state, but whether this reflects functional coupling or shared modulation is not always clear [21, 76, 77]. These challenges highlight the need to interpret correlation-based metrics with caution. Future work may benefit from integrating directed models [36, 83, 84] (e.g., Granger causality, transfer entropy), perturbation methods [85, 86], or predictive frameworks [19, 41, 87]. For example, dynamic Bayesian network (DBN) modeling has been used to track state-dependent changes in inter-laminar connectivity in area V4 [10, 48], offering a directed approach that could complement our framework in distinguishing communication from shared modulation.

Additionally, applying such approaches across cortical and subcortical systems may reveal whether population coupling reflects a conserved computational principle or varies with regional specialization. For instance, associative areas may show more internally-driven coupling, whereas sensory regions may be dominated by stimulus-locked dynamics. Ultimately, combining state estimation [48, 61, 87], multi-region laminar recordings [9, 10, 88], and causal inference [79, 85, 89] will be crucial for identifying when population coupling reflects communication versus shared modulation, advancing our understanding of functional architecture across distributed neural circuits.

## METHODS

Two male rhesus macaques (Macaca mulatta) participated in this study (Monkey D: 6 years old; Monkey M: 8 years old). The animals were pair-housed under a 12-hour light/dark cycle, and their water intake was regulated during the experimental period. All procedures were approved by the Yale University Institutional Animal Care and Use Committee and conformed to NIH guidelines.

### Surgical procedures

Surgical procedures followed protocols described in previous studies [24, 42, 85, 86]. Low-profile titanium recording chambers were implanted in two macaques, providing access to area V1—bilaterally in Monkey M and in the right hemisphere of Monkey D. Chamber placement was guided by sulcal reconstructions derived from preoperative structural MRI scans. Following implantation, the native dura mater was surgically removed and replaced with a transparent silicone artificial dura (AD), which enabled direct visualization of V1 cortical sites for precise probe targeting. All procedures were approved by the Yale University Institutional Animal Care and Use Committee and conformed to NIH guidelines.

### Electrophysiology

Prior to recordings, 64-channel laminar probes (NeuroNexus Technologies; 2 shanks, 32 channels per shank; 70 μm inter-site spacing; 200 μm inter-shank spacing) were electroplated with PEDOT using nanoZ (White Matter LLC). At the start of each session, a probe was inserted into V1 using a titanium mounting stage secured to the chamber and positioned with an electronic micromanipulator (Narishige Inc.). Probe orientation was verified to be orthogonal to the cortical surface via surgical microscope (Leica Microsystems).

To penetrate the brain, the probe first passed through the artificial dura (AD), arachnoid, and pia at high speed (>100 μm/s). Once the tip entered the cortex, the rest of the probe was advanced slowly (2 μm/s). After full insertion, the probe was retracted slightly (2 μm/s) to relieve pressure without altering its cortical position.

Neural signals were acquired at 30 kHz using a 64-channel digital headstage and recorded with the RHD Recording System (Intan Technologies). Spike sorting was performed offline using Kilosort2, followed by manual curation in Phy [1, 88]. Axonal spikes—identified by waveforms with a peak preceding the trough—were excluded.

### Behavioral control and eye tracking

Behavioral tasks were controlled using NIMH MonkeyLogic [90]. Eye position and pupil diameter were recorded at 120 Hz using an infrared eye tracker (ETL-200, ISCAN Inc.) and fed into the control system. Visual stimuli were presented on a monitor 57 cm from the monkey (60 Hz refresh rate). Trials were aborted if gaze deviated beyond 1.2 dva (monkey D) or 1.5 dva (monkey M) from fixation. In separate spontaneous sessions, monkeys sat in darkness without visual stimuli. Eye tracking continued to monitor eye state, enabling detection of eyes-open, eyes-closed, and saccade periods.

### Receptive field mapping

We mapped receptive fields (RFs) by presenting Gabor patch stimuli (2–4 cycles/degree, 0.25–1 dva Gaussian half-width, 100% contrast) on a square grid covering the lower visual quadrant (bilaterally in monkey M, left hemisphere in monkey D), while the monkey fixated centrally. Grid spacing was optimized per session and ranged from 0.25–1 dva. On each frame, a Gabor stimulus was presented at a random grid location and orientation. For each recording channel, we computed local field potential (LFP) power 40–200 ms after stimulus onset at each location. These power values were smoothed using a Gaussian kernel (σ = 0.75 dva), and the peak of the averaged LFP power across channels was defined as the column’s RF center. Spatial RF maps for each channel were visualized as stacked contour plots across the two shanks.

### Current source density mapping

To identify laminar boundaries and assess the relative strength of input to the superficial layer across visual conditions, we performed current source density (CSD) analysis [46]. While the monkeys maintained central fixation, high-contrast (100% luminance) white annular stimuli were briefly flashed (32 ms) at the center of the receptive field. Local field potentials (LFPs) were averaged across trials and spatially smoothed with a Gaussian kernel (σ = 140 μm). The CSD was computed as the second spatial derivative of the LFP and interpolated at 7 μm intervals. The Input layer was identified by the earliest current sink, corresponding to feedforward input to layer IV. Channels located above and below this sink were designated as superficial and deep layers, respectively.

### Experimental tasks

While the monkeys maintained fixation at the center of the screen, visual stimulus arrays were presented for 100 ms, followed by a 200–250 ms inter-stimulus interval. Each array consisted of a stimulus at the receptive field (RF) center, presented either in isolation or paired with a flanker at different spatial locations [42]. The resulting stimulus configurations were not distinguished for the purpose of this study and were aggregated for analyzing stimulus-evoked activity (see below). Stimulus configurations were randomly interleaved and repeated 4–6 times per trial.

The central stimuli were sine Gabor patches (25% luminance contrast, 3.5 cycles/degree, 0.5–1.0° Gaussian half-width), presented at 6 evenly spaced orientations and 2 opposite phases. Flankers were matched to probes in all parameters except contrast (100% luminance). In flanked trials, flanker orientation was either the same as or orthogonal to the probe, with equal probability.

In separate sessions, spontaneous neural activity was recorded while monkeys sat in complete darkness for 10-40 minutes. No visual stimuli were presented, and animals were not engaged in any task.

For stimulus-evoked activity, recordings were collected across 22 sessions (14 in monkey M, 8 in monkey D), yielding 1123 units in monkey M, and 677 units in monkey D. For spontaneous activity, recordings were collected across 11 sessions (3 in monkey M, 8 in monkey D), yielding 333 units in monkey M, and 526 units in monkey D. Units include single units and multi-unit clusters in all three layers.

### Data preparation

We categorized the spike data into four conditions: stimulus-evoked activity (stimulus-on and stimulus-off) and spontaneous activity (eyes-open and eyes-closed).

For stimulus-evoked activity, spikes for each unit were counted in 100-ms trial windows, starting from the response onset (stimulus-on bins) and ending 30 ms before the response onset (stimulus-off bins), with the 30-ms gap ensuring minimal contamination by sensory input. Responses to all stimulus types (gratings with varying orientations) were pooled to avoid reflecting the orientation selectivity of individual neurons. Response onset was computed separately for each laminar layer. Units with a mean firing rate below 0.2 spikes/s during stimulus-off periods were excluded. Baseline firing rates were defined using the peri-stimulus time histogram (PSTH) from –30 to +30 ms relative to stimulus onset. The response onset time was defined as the point when the PSTH exceeded the 95% confidence interval (CI) of the baseline. Units with no significant response were also excluded. These steps were applied separately to each layer to ensure response alignment across layers.

For spontaneous activity, we excluded periods of saccades and investigated periods when the animals maintained fixation with eyes open or had their eyes closed. Spikes were counted in 500-ms sliding trial windows during each condition. Mean firing rates were computed, and units firing below 0.2 spikes/s were excluded.

Spike counts were z-scored within each condition to isolate trial-to-trial fluctuations by removing mean activity. For spontaneous activity, each 500-ms window was treated as a trial. We analyzed stimulus-evoked activity from 5 sessions in monkey D and 11 sessions in monkey M, and spontaneous activity from 6 sessions in D and 3 in M. Sessions were included only if they contained sufficient units (see below) in both Input and Superficial layers to form source–target population pairs, and if the number of trials met minimum thresholds (stimulus-evoked: ≥3000; spontaneous: ≥360).

### Reduced-Rank Regression (RRR)

To quantify the low-dimensional predictive relationship between two neuronal populations, we applied a modified version of the approach described in [41, 42]. To mitigate overfitting due to noise (Fig. S1), we fit a Ridge regression model for *Y* = *XB*, where *X* and *Y* represent trial-to-trial fluctuations in activity from the source and target populations, respectively.

We then applied Reduced-Rank Regression (RRR), as previously described. This method performs principal component analysis (PCA) on the initial prediction *Y*, retaining components up to a specified rank. The optimal dimensionality was defined as the smallest number of dimensions that achieved prediction accuracy within one standard error of the mean (SEM) of the maximum (across cross-validation folds). We constructed a rank-reduced regressor *B*_*RRR*_ corresponding to the optimal predictive subspace. Prediction accuracy was quantified as 1 − *NSE*, where *NSE* is the normalized squared error, and computed across multiple resampled trial sets for each source–target pair.

For each recording session in either monkey, we randomly sampled *p* source neurons and *q* target neurons for regression. Sample sizes were scaled to the total number of available units per session to maintain comparable distributions of accuracy across sessions. Specifically, for monkey D: stimulus-evoked analysis used *p* = 15, *q* = 3 ; spontaneous analysis used *p* = 10, *q* = 3. For monkey M: stimulus-evoked analysis used *p* = 15, *q* = 3; spontaneous analysis used *p* = 30, *q* = 3.

To reduce the impact of within-session variability and noise, we performed multiple random draws of trials for each source–target pair. Final prediction accuracy was reported as the SEM of all positive accuracy values across resampling. Accuracy values less than zero—indicating predictions worse than the mean—were discarded, as they suggest no meaningful correlation between the sampled populations. A pair of populations was included in subsequent analysis only if both compared conditions yielded a positive accuracy.

### Leading principal angle (LPA)

To compare the predictive subspaces of the same source–target neuron pair across different conditions, we computed the leading principal angle (LPA) between the predictive subspace *B*_*RRR*_ obtained under each condition. For a given neuron pair, the predictive subspaces under two conditions were denoted *B*_1_ and *B*_2_. The principal angles*θ*_1_, *θ*_2_, … between these subspaces were derived from the singular values of the matrix 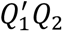, where *Q*_1_ and *Q*_2_ are the orthonormal bases of *B*_1_ and *B*_2_, respectively. The LPA *θ*_1_ corresponds to the smallest angle between any two vectors in the subspaces *B*_1_ and *B*_2_, and cos *θ*_1_ quantifies the maximal similarity between the predictive directions under the two conditions. Importantly, this method does not require *B*_1_ and *B*_2_ to have the same dimensionality, allowing us to compare predictive subspaces even when the optimal ranks differ across conditions.

LPA values were included in the final analysis only when both conditions yielded positive prediction accuracy for the same sampled source–target pair. To assess the reliability of LPA estimates, we also computed the within-condition variance, defined as the LPA between subspaces obtained from non-overlapping resampled trials within the same condition. Finally, to establish a baseline, we computed the LPA between random subspaces of the same dimensionality as *B*_1_ and *B*_2_, allowing for comparison against chance-level alignment.

### Fractional optimal dimension

For each source–target neuron pair, we computed the fractional optimal dimension, defined as the optimal rank obtained from RRR divided by the maximum possible dimension (the minimum of the number of source and target units). Only pairs with positive prediction accuracy were included in this analysis, and the fractions were summarized using estimation statistics of the mean.

### Modulation index (MI)

We quantified changes in prediction accuracy across conditions using a modulation index (MI), computed for each source–target pair as:

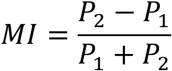

where *P*_1_ and *P*_2_ denote prediction accuracies under two different conditions. MI values were reported with estimation statistics of the mean, reflecting the magnitude and direction of modulation in inter-population predictability.

### Delay analysis

To examine the directionality of inter-layer interactions, we performed a delay analysis by predicting superficial-layer activity from input-layer activity at varying temporal offsets. While prior results were based on simultaneous recordings, this analysis introduced a temporal shift in the superficial activity from −10 ms to +25 ms relative to the input activity. The analysis pipeline of reduced-rank regression and accuracy computation remained unchanged. For each time lag, prediction accuracy was computed using the same cross-validated procedure and summarized with estimation statistics across source-target pairs and draws of trials.

### Recurrent neural network (RNN) simulation

We implemented a recurrent neural network consisting of 200 units, organized into two layers: an Input layer and a Superficial layer, each containing 100 units. Within each layer, the recurrent connectivity matrices *J*_*I*–*I*_ and *J*_*S*–*S*_ were defined such that all off-diagonal weights *w*_*ij*_ = 0.001 for *i* ≠ *j*; self-connections were not included. To simulate a low-rank, feedforward projection from the Input to the Superficial layer, we generated 30 pairs of random vectors *m*_1×100_ and *n*_1×100_, where each *m*_*i*_ ∼ 𝒩(2, 0.8^2^) and *n*_*i*_ ∼ 𝒩(1, 0.4^2^). The inter-layer connectivity matrix was then defined as 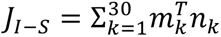, resulting in a rank-30, strictly positive matrix. No feedback was implemented with *J*_*S*–*I*_ = 0. We simulated three conditions:

**1. Eyes-open/Stimulus-off**: baseline input *I*_1_ (*t*) only

*I*_1_ (*t*) was modeled as Gaussian noises *η*_*i*_ (*t*)∼𝒩(0, 0.01^2^) sampled independently for each of the 200 units from a standard normal distribution:

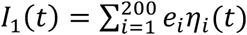

where *e*_*i*_ is the basis vector of neuron *i*. Independent noises were updated at 10 Hz (relative to the network’s time constant; not tied to real-world time). The mean firing rate remains near 0, serving as a baseline of the network activity.

**2. Stimulus-on**: *I*_1_ (*t*) + sensory input *I*_2_ (*t*)

*I*_2_ (*t*) targeted the 100 Input-layer units only. It consisted of 30 unique patterns, each a vector *v*_*i*_ of size 100, sampled from *v*_*i*_ ∼ 𝒩(0. 1, 0.1^2^).

The positive mean pattern ensured net positive drive through the feedforward weights to the Superficial layer. To simulate the temporal co-fluctuation, these patterns were injected as square waves *f*_*d*_ (*t*):

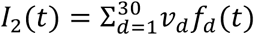

where each *f*_*d*_ (*t*) has random amplitudes sampled from 𝒩(0.1, 0.003^2^) with a periodicity of 40 s. The positive mean amplitude ensures a time-averaged positive input, resulting in an elevated overall network firing rate, consistent with observations in real neurons.

**3. Eyes-closed**: *I*_1_ (*t*) + wakefulness-related input *I*_3_ (*t*)

*I*_3_ (*t*) targeted all 200 units simultaneously. It consisted of 30 unique patterns 𝑢_*i*_ ∼ 𝒩(0, 0.1^2^) of size 200, where zero mean pattern ensures that inter-layer communication was minimally engaged. These patterns were also delivered as square waves with random amplitudes at a periodicity of 40 s:

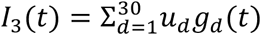

where each *f*_*d*_ (*t*) has random amplitudes sampled from 𝒩(0, 0.018^2^). The mean amplitude is set to zero to reflect that wakefulness does not produce a similar elevation in mean firing rate as the sensory input.

### Regression on simulated data

The network was initialized with random states and run for 3200 s. Spike-like activity was binned using a 10-s window with a 0.5-s step size. After excluding an initial transient period, over 4000 trials remained for analysis across all three conditions. For each of the three conditions, we applied reduced-rank regression by randomly sampling 20 source and 20 target units and drawing 1,000 trials per iteration. Each iteration used 10-fold cross-validation. The same analysis was conducted for both the directed and undirected models of the Input and Superficial populations. We also repeated the LPA and MI analyses, where the introduction of *I*_2_ reflects the effect of visual stimulation, and the introduction of *I*_3_ captures the effect of the eyes-closed state relative to the eyes-open condition.

### Directionality and activity structure

To investigate the directionality of modulation between layers in the simulated network, we manipulated the relative amplitudes of Gaussian noise *I*_1_ across the Input and Superficial layers. By varying noise amplitudes, we effectively altered the degree to which the structured patterns *I*_2_ or *I*_3_ emerged in each layer and contributed to shared co-fluctuations across populations.

To quantify the level of structure in a population’s activity, we analyzed the dominance of its principal components. In a population driven purely by random noise, principal components contribute more evenly to the variance, producing a near-linear increase in the cumulative variance explained across component order. In contrast, structured activity tends to be dominated by a few leading components, indicating a low-dimensional subspace. We computed the variance explained by each principal component using singular value decomposition (SVD) on z-scored population activity. As a summary measure of structure, we defined the area under the cumulative variance explained curve (AUC). A larger AUC indicates a stronger concentration of variance in the top components, reflecting greater activity structure and lower effective dimensionality.

## AUTHOR CONTRIBUTIONS

YX and MPJ conceptualized the project. MPM collected the electrophysiological data, and ASN supervised data collection. YX analyzed the electrophysiological data and performed computational modeling. MPJ supervised the project. YX, ASN, and MPJ wrote the manuscript.

## ACKNOWLEDGEMENTS

This research was supported by NIH R01 EY034605, NIH R00 EY025026, NIH R21 MH126072 and SFARI 875855 to MPJ, NARSAD Young Investigator Grant, Ziegler Foundation Grant, Yale Orthwein Scholar Funds, NIH R01 EY032555, NIH R21 MH126072 and SFARI 875855 to ASN, and by NEI core grant for vision research P30 EY026878 to Yale University. We thank the veterinary and husbandry staff at Yale for excellent animal care.

## SUPPLEMENTARY FIGURES

**Fig. S1.**
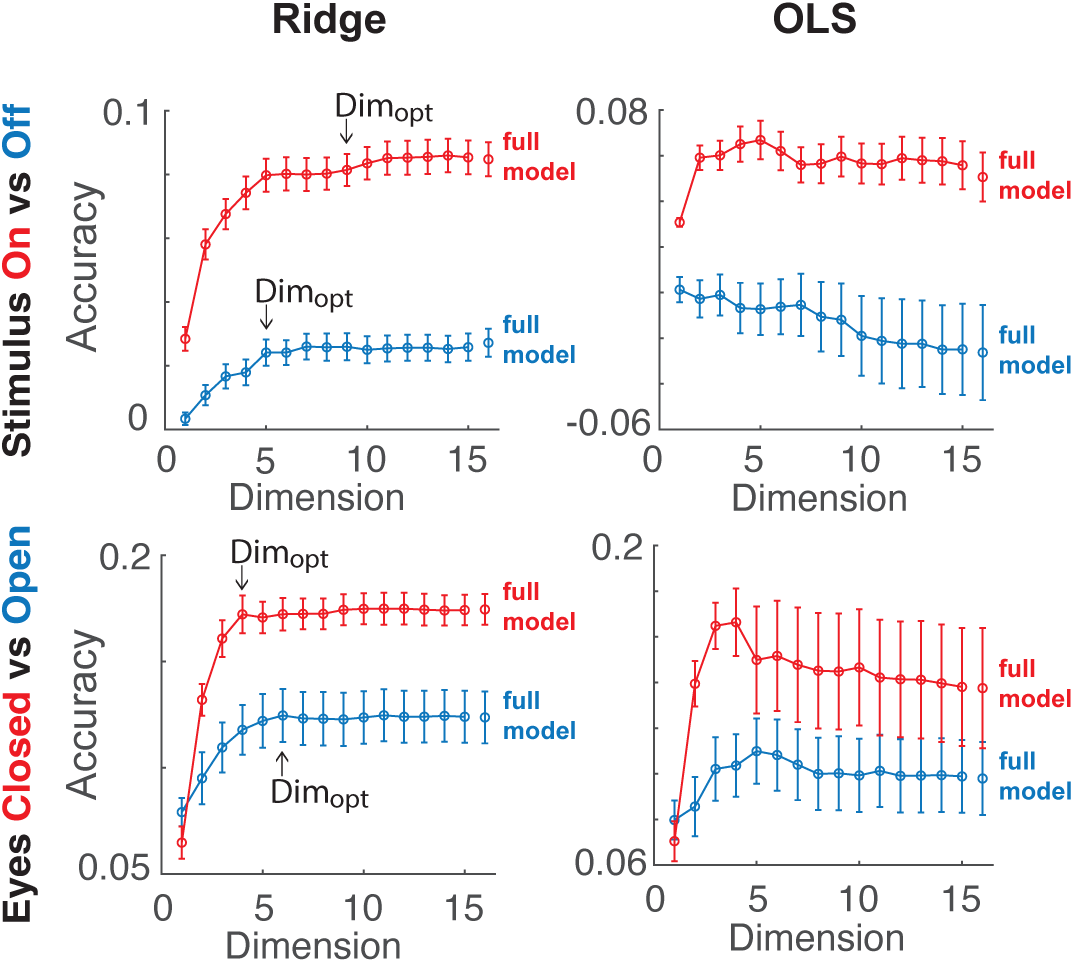
Ridge reduced-rank regression vs. OLS reduced-rank regression. Example sessions from monkey D illustrating how overfitting was mitigated during dimensionality increases in reduced-rank regression by applying Ridge regression in place of ordinary least squares (OLS). Predictions were based on 32 source and 20 target units for stimulus-on vs. stimulus-off, and 24 source and 20 target units for eyes-closed vs. eyes-open conditions. Error bars denote the standard error of prediction accuracy, computed across multiple trial resamples and corresponding cross-validation folds.

**Fig. S2.**
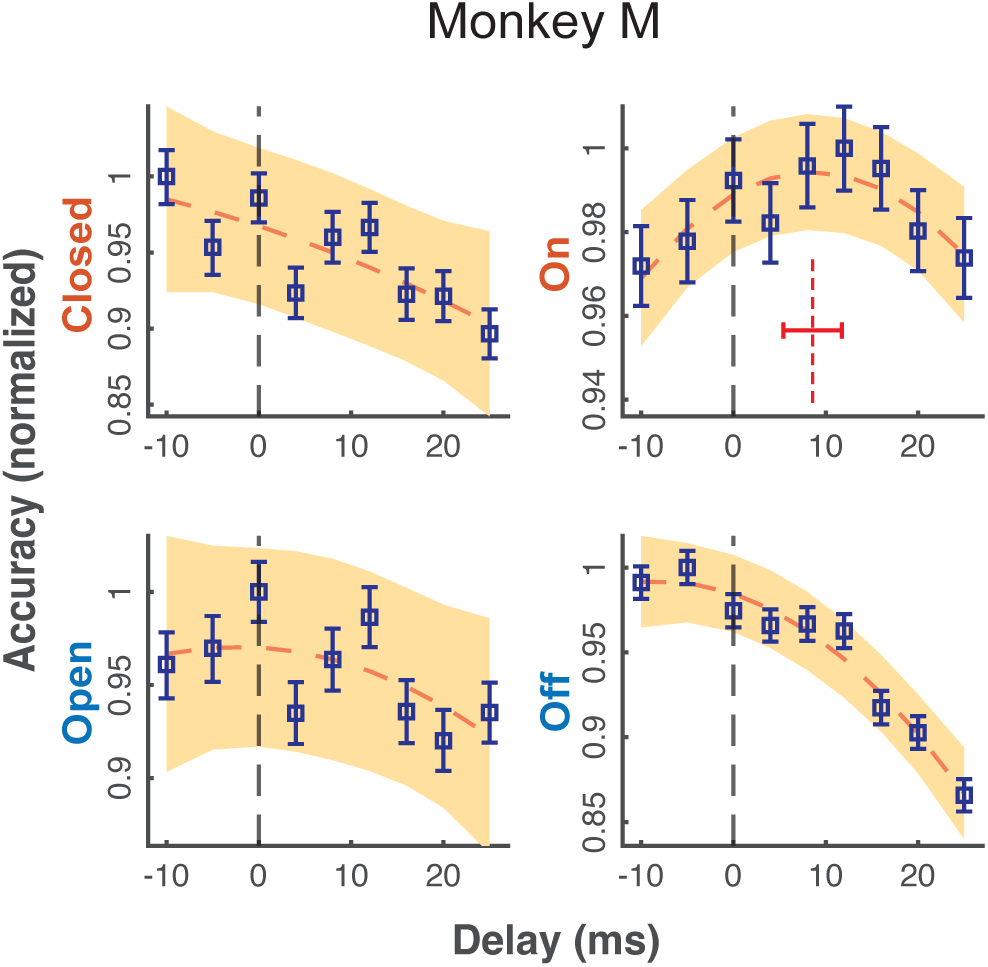
Delay analysis of the directed model in monkey M. Same as Fig. 2C. The target activity of the superficial population is shifted in the range -10 ms to 25 ms w.r.t. input layer activity. The accuracy is normalized, then fitted to a quadratic function of the delay time. Yellow areas represent the 95% CIs of the normalized efficacy. The vertical red dashed line in stimulus-on condition shows the symmetry axis (optimal delay), along with the standard error.

**Fig. S3.**
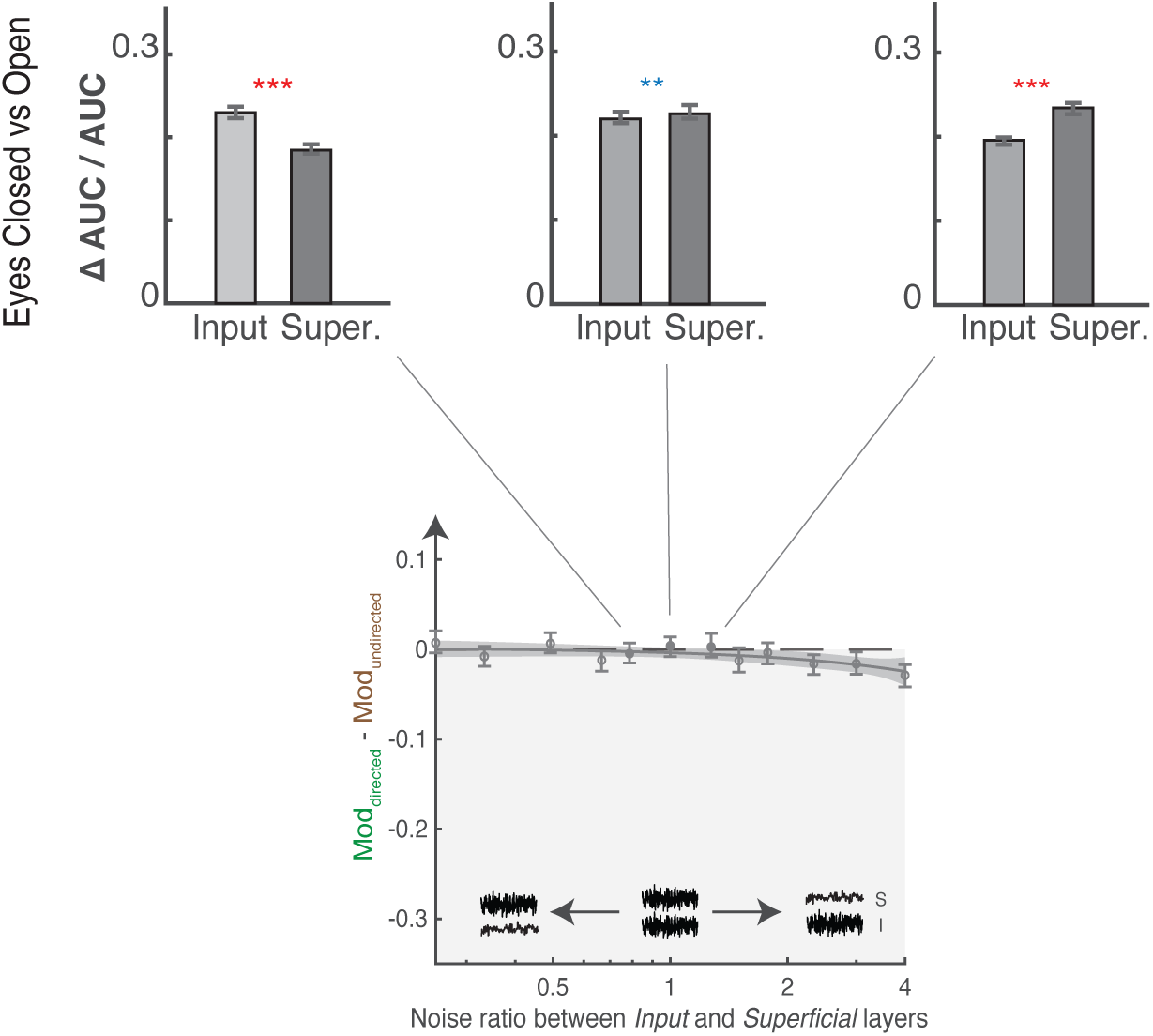
Relative activity structure for spontaneous modulation with different parameters. Top panel shows the increase of structured activity in input and superficial layers under eyes closed vs. open conditions. Error bars indicate the 95% CIs of the mean. From left to right, results were plotted for Noise ratio = 0.79 (Fig. 4F), 1, 1.27, associated with the solid circles in the bottom panel (Fig. 4B). Noise ratio = 0.79 best captures the empirical findings (Fig. 4G).

